# Recycling of single-stranded DNA-binding protein by the bacterial replisome

**DOI:** 10.1101/486555

**Authors:** Lisanne M. Spenkelink, Jacob S. Lewis, Slobodan Jergic, Zhi-Qiang Xu, Andrew Robinson, Nicholas E. Dixon, Antoine M. van Oijen

## Abstract

Single-stranded DNA-binding proteins (SSBs) support DNA replication by protecting single-stranded DNA from nucleolytic attack, preventing intra-strand pairing events, and playing many other regulatory roles within the replisome. Recent developments in single-molecule approaches have led to a revised picture of the replisome that is much more complex in how it retains or recycles protein components. Here we visualise how an *in vitro* reconstituted *E. coli* replisome recruits SSB by relying on two different molecular mechanisms. Not only does it recruit new SSB molecules from solution to coat newly formed single-stranded DNA on the lagging strand, but it also internally recycles SSB from one Okazaki fragment to the next. We show that this internal transfer mechanism is balanced against recruitment from solution in a manner that is concentration dependent. By visualising SSB dynamics in live cells, we show that both internal transfer and external exchange mechanisms are physiologically relevant.

## INTRODUCTION

The majority of processes associated with DNA metabolism involve the generation of single-stranded DNA (ssDNA). As a transient species that ultimately needs to be reconverted into more stable doublestranded DNA (dsDNA), ssDNA acts as a substrate for a large number of pathways. A key protein in the initial steps of ssDNA processing is the ssDNA-binding protein (SSB), which coats naked ssDNA and thus protects it from nucleolytic attack and prevents intra-strand pairing events such as hairpin formation. Further, it plays a critical role in the organisation of protein–protein and protein–DNA interactions within the replisome, the protein machinery responsible for DNA replication (1–4).

*Escherichia coli* SSB is a stable homotetramer with each 177 amino acid-subunit separated into two distinct domains (5). The N-terminal domain (112 residues) forms an oligonucleotide/oligosaccharide-binding (OB) fold responsible for ssDNA binding (6). The C-terminal domain is more variable, except for a highly conserved acidic C-terminal tail, which serves as an interaction site for many binding partners (2,4,7,8). SSB can bind to ssDNA in different modes depending on the concentration of cations and the SSB/ssDNA stoichiometry (9). The prevalent binding modes observed in *in vitro* studies are the SSB_65_ and SSB_35_ modes, corresponding to the binding of 65 and 35 nucleotides to the SSB tetramer, respectively (10). In the SSB_65_ mode, favoured at moderately high salt concentrations (11), all four ssDNA-binding sites are bound to ssDNA (Figure 1B, left). In the SSB_35_ binding mode, favoured in low salt concentrations (12), only two ssDNA-binding sites are occupied (Figure 1B, right) (6); the negative cooperativity that restricts occupancy to two sites in the SSB_35_ mode provides a facile mechanism for transfer of SSB from one ssDNA molecule to another without its dissociation into solution (10,13,14).

**Figure 1.**
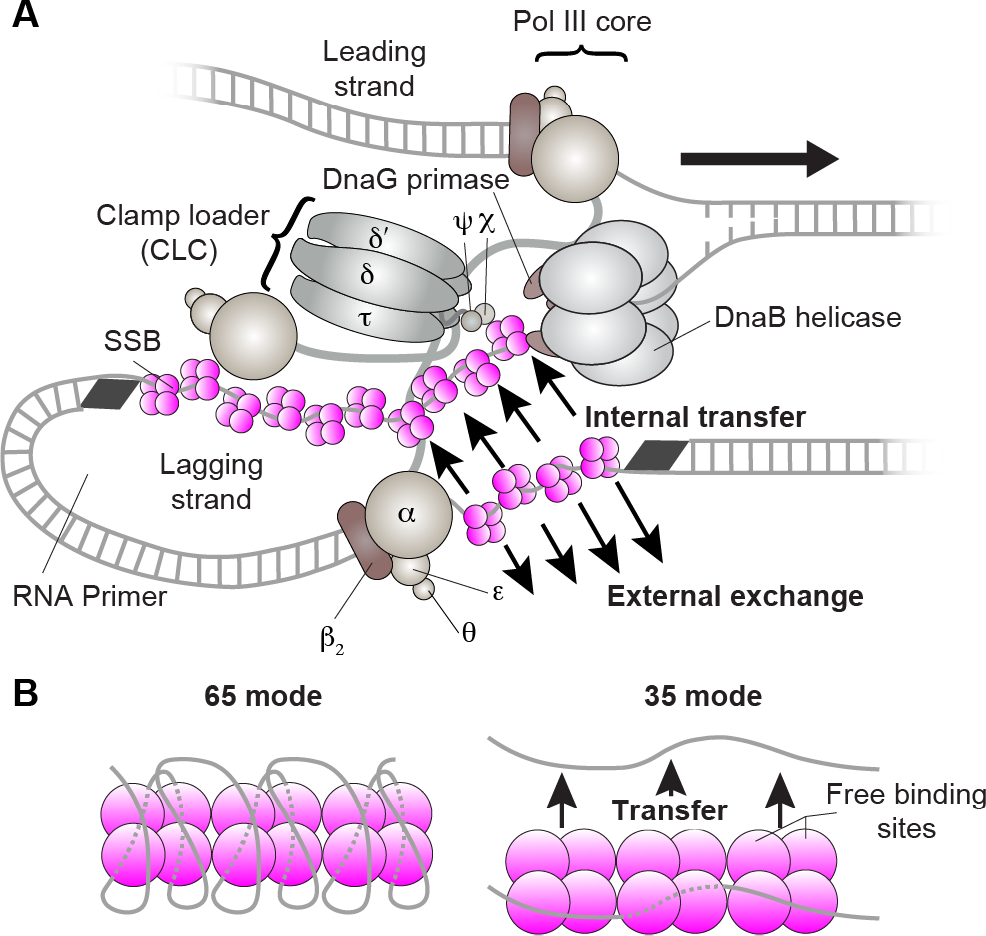
The *E. coli* replisome. (**A**) Schematic representation of the organisation of the *E. coli* replication fork. The DnaB helicase encircles the lagging strand, facilitates unwinding of dsDNA through ATP hydrolysis, and recruits DnaG primase for synthesis of RNA primers that initiate synthesis of 1–2 kb Okazaki fragments on the lagging strand. The Pol III holoenzyme (HE) uses the ssDNA of both strands as a template for simultaneous synthesis of a pair of new DNA duplex molecules. The β_2_ sliding clamp confers high processivity on the Pol III HE by tethering the αεθ Pol II core onto the DNA. The clamp loader complex (CLC) assembles the β_2_ clamp onto RNA primer junctions. Up to three Pol III cores interact with the CLC through its τ subunits to form the Pol III* complex, with the τ subunits also binding to DnaB, thus coupling the Pol III HE to the helicase. The ssDNA extruded from the DnaB helicase is protected by SSB (33,67). (**B**) Different DNA-binding modes of SSB. In the SSB_65_ mode, all four OB domains are bound to DNA (left). In the SSB_35_ mode, only two DNA-binding sites are occupied. The observation of transfer of SSB between discrete ssDNAs in this mode suggests a possible internal-transfer mechanism.

During DNA replication, ssDNA is produced when the helicase unwinds the parental dsDNA. On one of the daughter strands, the leading strand, new DNA is synthesised continuously by a copy of the DNA polymerase III (Pol III) core closely tracking and travelling in the same direction as the helicase (Figure 1A), thereby minimising the amount of exposed ssDNA. Indeed, uncoupled leading-strand replication occurs efficiently in the complete absence of SSB (15). On the other (lagging) strand, DNA is synthesised discontinuously. Due to the opposite polarities of the two DNA strands and the ability of polymerases to extend a DNA chain only in the 3′ to 5′ direction, the Pol III core on the lagging strand synthesises DNA in the direction opposite to that of the moving fork (16,17). As a result, stretches of ssDNA are generated on the lagging strand that are not converted into dsDNA until the next Okazaki fragment is primed and synthesised. During the period these stretches of ssDNA are exposed, they are coated with SSB. As new DNA is synthesised on the lagging strand, SSB has to be displaced by the advancing polymerase, perhaps involving an interaction of its C-terminal tail with the χ subunit of the Pol III holoenzyme complex (18,19).

Biochemical studies of SSB in the last decades have used short oligonucleotides, outside the context of active replisomes, leading to two different models of the dynamics of its binding to and dissociation from ssDNA within the replication complex. In the first model, newly exposed ssDNA is bound by SSB from the cytosol. It has been shown that SSB binds to free ssDNA in a diffusion-controlled process (13,20). With the estimated *in vivo* SSB concentration of 300–600 nM (5,21–24), such rapid binding would lead to efficient coating of newly exposed ssDNA within milliseconds. In this model, subsequent displacement of SSB during filling in of the gap by the lagging-strand Pol III core will cause the SSB to diffuse back into the cytosol.

In the second model, SSB is effectively recycled within the replisome through an internal-transfer mechanism. Studies using surface plasmon resonance and nano-electrospray ionisation mass spectrometry showed that the SSB_35_ mode supports transfer of SSB tetramers between discrete oligonucleotides (14). Stopped-flow experiments demonstrate similar behaviour on rapid time scales, suggesting that transfer occurs without proceeding through a protein intermediate that is free from DNA (13). Instead, transfer involves a transiently paired intermediate during which SSB is ‘handed’ from the first to the second ssDNA with the system going through a state in which the tetramer is bound to two strands simultaneously (Figure 1B, right).

The biochemical studies have been unable to directly visualise the dynamic behaviour of SSB within the replisome. As a result, it is unknown how the replication machinery recruits SSB and whether it may retain it during multiple cycles of Okazaki-fragment synthesis. It is unclear whether the approximately 500 copies of SSB available within the cell are sufficient to support rapid coating of all ssDNA during fast growth, with up to 12 replisomes active simultaneously (25), or whether internal recycling mechanisms are operative that enable a replisome to maintain its own local pool of SSB molecules.

Here, we directly probe the existence of a process in which transfer of SSB occurs from in front of the Pol III to the newly-exposed ssDNA behind the helicase on the lagging strand, without its dissociation into the cytosol (Figure 1A). To access this mechanism experimentally, we use single-molecule fluorescence imaging to visualise the dynamics of SSB during active DNA replication, both *in vitro* in a reconstituted replication reaction and inside living bacterial cells. We rely on the strength of the single-molecule approach to visualise transient intermediates and acquire detailed kinetic information that would otherwise be hidden by the averaging inherent to ensemble measurements (26–29). Particularly, we show that SSB is recycled within the replisome on time scales corresponding to the synthesis of multiple Okazaki fragments, verifying the existence of an internal-transfer mechanism. At higher SSB concentrations, however, we see that this mechanism competes with external exchange to and from solution. Using *in vivo* single-molecule imaging, we show that both processes occur at the replication fork. Our observations suggest that the interactions controlling association and dissociation of SSB within the replisome provide a balance between plasticity and stability, enabling its exchange when available, but ensuring replisome stability in its absence from the cellular environment.

## MATERIALS AND METHODS

### Replication proteins

*E. coli* DNA replication proteins were produced from *E. coli* strains with genes from *E. coli* MG1655 as described previously: the β_2_ sliding clamp (30); SSB (14); the DnaB_6_(DnaC)_6_ helicase–loader complex (31); DnaG primase (32); the Pol III τ_3_δδ′ψχ clamp loader (15); and Pol III αεθ core (33).

### Expression and purification of SSB-K43C

Plasmid pND539 was constructed by insertion of a *Bam*HI–*Eco*RI fragment of pND73 (14) between the same sites in the λ-promoter phagemid vector pMA200U (34). Oligonucleotide-directed mutagenesis with single-stranded pND539 was then used to introduce a cysteine codon in place of the lysine-43 codon of *ssb* to yield phagemid pCL547; the mutation was confirmed by nucleotide sequence determination. Expression and purification of the single cysteine mutant of SSB, SSB-K43C were carried out as previously described for wild-type SSB (14).

### Labelling of SSB-K43C

Methods described below were adapted from Kim *et al.* (35). Three different fluorescent probes were used to label SSB-K43C: Alexa Fluor 488, 555, and 647 (Invitrogen). First, a total of 6.3 mg of SSB-K43C was reduced with 3 mM tris(2-carboxyethyl)phosphine (pH 7.6) in precipitation buffer (100 mM sodium phosphate pH 7.3, 200 mM NaCl, 1 mM EDTA, 70% (*w/v*) ammonium sulphate) at 6°C for 1 h with gentle rotation to yield Fraction I. Fraction I was centrifuged (21,000 × *g*; 15 min) at 6°C and the supernatant carefully removed. The precipitate was washed with ice cold precipitation buffer that had been extensively degassed by sonication and deoxygenated using Ar gas, then pelleted by centrifugation (21,000 × *g*; 15 min) at 6°C and supernatant removed to yield Fraction III. The labelling reaction was carried out on Fraction III, now devoid of reducing agent, using 40 μM of maleimide-conjugated dyes with 84 μM SSB-K43C in 500 μl of deoxygenated and degassed buffer (100 mM sodium phosphate pH 7.3, 200 mM NaCl, 1 mM EDTA). The reaction was allowed to proceed for 3 h at 23°C in the dark. The reaction was subsequently quenched using 30 mM dithiothreitol for 1 h at 6°C, yielding Fraction IV. Fraction IV was applied at 1 ml min^−1^ to a column (1.5 × 10 cm) of Superdex G-25 (GE-Healthcare) resin equilibrated with gel filtration buffer (50 mM Tris.HCl pH 7.6, 3 mM dithiothreitol, 1 mM EDTA, 100 mM NaCl). Fractions containing the labelled SSB-K43C were pooled and dialysed into storage buffer (50 mM Tris.HCl pH 7.6, 3 mM dithiothreitol, 1 mM EDTA, 100 mM NaCl, 20% (*v/v*) glycerol). The degree of labelling was determined by UV/Vis spectroscopy to be between 1 and 2 fluorescent dyes per SSB tetramer.

### Single-molecule rolling-circle assay

Construction of the 2030-bp template used for most rolling-circle assays has been described (36). To construct the M13 rolling circle template (37), the 66-mer 5′-biotin-T_36_AATTCGTAATCATGGTCATAGCTGTTTCCT-3′ (Integrated DNA Technologies) was annealed to M13mp18 ssDNA (New England Biolabs) in TBS buffer (40 mM Tris-HCl pH 7.5, 10 mM MgCl_2_, 50 mM NaCl) at 65°C. The primed M13 was then extended by adding 64 nM T7 gp5 polymerase (New England Biolabs) in 40 mM Tris-HCl pH 7.6, 50 mM potassium glutamate, 10 mM MgCl_2_, 0.1 mg ml^−1^ BSA, 5 mM dithiothreitol and 600 μM dCTP, dGTP, dATP and dTTP at 37°C for 60 min. The reaction was quenched with 100 mM EDTA and the DNA was purified using a PCR purification kit (Qiagen).

Microfluidic flow cells were prepared as described (38). Briefly, a PDMS flow chamber was placed on top of a PEG-biotin-functionalised microscope coverslip. To help prevent non-specific interactions of proteins and DNA with the surface, the chamber was blocked with buffer containing 20 mM Tris-HCl pH 7.5, 2 mM EDTA, 50 mM NaCl, 0.2 mg ml^−1^ BSA, and 0.005% Tween-20. The chamber was placed on an inverted microscope (Nikon Eclipse Ti-E) with a CFIApo TIRF 100× oil-immersion TIRF objective (NA 1.49, Nikon) and connected to a syringe pump (Adelab Scientific) for flow of buffer.

Conditions for coupled DNA replication under continuous presence of all proteins were adapted from previously described methods (15,33,37). All *in vitro* single-molecule experiments were performed at least four times. Briefly, 30 nM DnaB_6_(DnaC)_6_ was incubated with 1.5 nM biotinylated ds M13 template in replication buffer (25 mM Tris-HCl pH 7.9, 50 mM potassium glutamate, 10 mM Mg(OAc)_2_, 40 μg ml^−1^ BSA, 0.1 mM EDTA and 5 mM dithiothreitol) with 1 mM ATP at 37°C for 30 s. This mixture was loaded into the flow cell at 100 μl min^−1^ for 40 s and then at 10 μl min^−1^. An imaging buffer was made with 1 mM UV-aged Trolox, 0.8% (*w/v*) glucose, 0.12 mg ml^−1^ glucose oxidase, and 0.012 mg ml^−1^ catalase (to increase the lifetime of the fluorophores and reduce blinking), 1 mM ATP, 250 μM CTP, GTP and UTP, and 50 μM dCTP, dGTP, dATP and dTTP in replication buffer. Pol III* was assembled *in situ* by incubating τ_3_δδ′ψχ (410 nM) and Pol III cores (1.2 μM) in imaging buffer at 37°C for 90 s. Replication was initiated by flowing in the imaging buffer containing 6.7 nM Pol III*, 30 nM β_2_, 300 nM DnaG, 30 nM DnaB_6_(DnaC)_6_ and SSB_4_ where specified at 10 μl min^−1^. Reactions were carried out at 31°C, maintained by an electrically heated chamber (Okolab).

Double-stranded DNA was visualised in real time by staining it with 150 nM SYTOX Orange (Invitrogen) excited by a 568-nm laser (Coherent, Sapphire 568-200 CW) at 150 μW cm^−2^. The red labelled SSB was excited at 700 μW cm^−2^ with a 647 nm laser (Coherent, Obis 647-100 CW). For simultaneous imaging of DNA and SSB, the signals were separated *via* dichroic mirrors and appropriate filter sets (Chroma). Imaging was done with an EMCCD camera (Photometics, Evolve 512 Delta). The analysis was done with ImageJ using in-house built plugins. The rate of replication of a single molecule was obtained from its trajectory and calculated for each segment that has constant slope.

Conditions for the pre-assembly replication reactions for the Okazaki fragment length measurements were adapted from published methods (33,39,40). Solution 1 was prepared as 30 nM DnaB_6_(DnaC)_6_, 1.5 nM biotinylated ds M13 substrate and 1 mM ATP in replication buffer. This was incubated at 37°C for 3 min. Solution 2 contained 60 μM dCTP and dGTP, 6.7 nM Pol III*, and 74 nM β_2_ in replication buffer (without dATP and dTTP). Solution 2 was added to an equal volume of solution 1 and incubated for 6 min at 37°C. This was then loaded onto the flow cell at 100 μl min^−1^ for 1 min and then 10 μl min^−1^ for 10 min. The flow cell was washed with replication buffer containing 60 μM dCTP and dGTP. Replication was finally initiated by flowing in the imaging buffer containing 50 nM β_2_, 300 nM DnaG and SSB_4_ where specified at 10 μl min^−1^.

Conditions for the chase replication reactions omitting SSB from solution during replication were set up as a normal continuous flow experiment. Reactions were allowed to proceed for 1 min before a replication mixture omitting only SSB was loaded at 10 μl min^−1^. To visualise the behaviour of SSB at a concentration of 100 nM, a 1:5 mixture of labelled and unlabelled SSB K43C was used.

To obtain the characteristic exchange time τ from the FRAP experiments, the data were fit with a FRAP recovery function, corrected for photobleaching (Equation 1, where *a* is the amplitude of photobleaching, τ_b_ is the photobleaching time, and I_0_ is the number of SSB molecules at the fork at steady state):

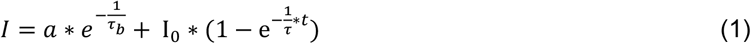

### Ensemble Okazaki-fragment length measurements

Coupled leading-and lagging-strand DNA synthesis reactions were set up in replication buffer (25 mM Tris-HCl pH 7.9, 50 mM potassium glutamate, 10 mM Mg(OAc)_2_, 40 μg ml^−1^ BSA, 0.1 mM EDTA and 5 mM dithiothreitol) and contained 1.0-1.5 nM 5′-biotinylated flap-primed 2-kb circular dsDNA template, 1 mM ATP, 250 μM CTP, GTP and UTP, and 50 μM dCTP, dGTP, dATP and dTTP, 6.7 nM Pol III*, 30 nM β_2_, 300 nM DnaG, 30 nM DnaB_6_(DnaC)_6_, and SSB_4_ as specified, in a final volume of 12 μl. Components (except DNA) were mixed and treated at room temperature, then cooled in ice for 5 min before addition of DNA. Reactions were initiated at 30°C, and quenched after 30 min by addition of 7 μl 0.5 M EDTA and 6 μl DNA loading dye (6 mM EDTA, 300 mM NaOH, 0.25% (*v/v*) bromocresol green, 0.25% (*v/v*) xylene cyanol FF, 30% (*v/v*) glycerol). The quenched mixtures were loaded into a 0. 6% (*w/v*) agarose gel in alkaline running buffer (50 mM NaOH, 1 mM EDTA). Products were separated by agarose gel electrophoresis at 14 V for 14 h. The gel was then neutralised in 1 M Tris-HCl, pH 7.6, 1.5 M NaCl and stained with SYBR Gold. The Okazaki fragment length distribution was calculated by normalising the intensity as a function of DNA length.

### *E. coli* strains with fluorescent chromosomal fusions

The strain EAW192 (*dnaQ-mKate2*) encodes a fusion of *dnaQ* with *mKate2* (33). JJC5380 (*ssb-YPet*) is MG1655 *ssb-YPet* Kan^R^ obtained by P1 co-transduction of the *ssb-YPet* fusion with the adjacent Kan^R^ marker from the AB1157 *ssb-YPet* Kan^R^ strain (41), and was a gift from Bénédicte Michel. The two-colour strain LMS001 (*ssb-YPet*, *dnaQ-mKate2*) was constructed by P1 transduction; JJC5380 cells (*ssb-YPet*) were infected with P1 grown on EAW192 (*dnaQ-mKate2*). Transductants were selected for kanamycin resistance. The strain JJC5945 (*dnaX-YPet*) is MG1655 *dnaX-YPet* (42). Wild-type (MG1655) and *dnaQ-mKate2* (EAW192) *E. coli* cells were cultured in LB. The *ssb-YPet* (JJC5380), *dnaQ-mKate2*, *ssb-YPet* (LMS001) and *dnaX-YPet* (JJC5945) strains were grown in LB supplemented with 25 μg/ml kanamycin.

### Growth rates of strains with fluorescent chromosomal fusions

To verify that the C-terminal labelling of SSB does not affect cell growth, we compared growth rates of five *E. coli* strains. We compared wild-type *E. coli* with *dnaQ-mKate2*, *ssb-YPet*, and the doubly labelled *dnaQ-mKate2* + *ssb-YPet* strains. We added the *dnaX-YPet* strain as a control. Single colonies of wild-type *E. coli* MG1655 and derivatives containing the C-terminal chromosomal *dnaX*, *dnaQ* and *ssb* fusions were used to inoculate 5 ml of L broth (with 25 μg ml^−1^ kanamycin, if required) and grown at 37°C with shaking overnight. L broth (100 ml) was inoculated with 1.0 × 10^5^ cells/ml from overnight cultures. Subsequent growth of each strain was monitored at 37°C on a plate reader (POLARstar Omega, BMG Labtech) determining OD_700_ every 20 min for 10 h. The labelled *ssb-YPet* and *dnaQ-mKate2* cells have similar growth rates to wild-type cells (Supplementary Figure S1A), indicating that labelling the SSB and DnaQ components of the replisome does not significantly disrupt DNA replication.

### *In vivo* FRAP measurements

The cells were grown at 37°C in EZ rich defined medium (Teknova) that included 0.2% (*w/v*) glucose. For imaging, cells were immobilised on coverslips that were functionalised with 3-aminopropyl triethoxysilane (Sigma Aldrich) (42) and then placed on the heated stage (Pecon) of the microscope (Olympus IX81, equipped with UAPON 100XOTIRF). Imaging was done at 37°C. FRAP measurements were performed using an automated fast filter wheel (Olympus U-FFWO) with a 50 μm pinhole in the back focal plane of the microscope. A 514 nm laser (Coherent, Sapphire 514-150 CW) was used for visualisation and photobleaching. FRAP pulses were 200 ms at 200 W cm^−2^ with the pinhole in place. Subsequent visualisation without the pinhole was done at 2 W cm^−2^. Imaging was done with an EMCCD camera (Hamamatsu c9100-13). The FRAP experiments were performed in triplicate, resulting in a total of 30 photobleached foci that were used for analysis. The image processing was done with ImageJ using in-house built plugins.

## RESULTS

### Visualisation of SSB *in vitro*

We use a fluorescence imaging approach to directly visualise DNA replication in real time to monitor the dynamics of SSB at the replication fork in a single-molecule rolling-circle assay, a method that provides information on the rate of production of new DNA by individual replisomes (37,43) while simultaneously enabling the visualisation of fluorescently labelled replisome components (33,44). A 5′-flap within a 2.0-kb double-stranded circular DNA substrate (36) is anchored to the surface of a microfluidic flow cell (Figure 2A). Replication is initiated by introducing a laminar flow of buffer containing the minimal set of 12 replication proteins required for coupled leading-and lagging-strand synthesis (Figure 1A). Replisomes assemble onto the fork structure within the circle and initiate unwinding and synthesis (15,37). As replication proceeds, the newly synthesised leading strand is displaced from the circle by helicase action to provide the template for lagging-strand synthesis. The net result of this process is the generation of a dsDNA tail that is stretched in the buffer flow and whose growth moves the tethered dsDNA circle away from the anchor point at a rate determined by the replication rate (Figure 2A). Replication is visualised by real-time near-total internal reflection fluorescence (TIRF) imaging of stained dsDNA (Figure 2B). Quantification of the instantaneous rates of individual replisomes resulted in an average single-molecule rate of 630 ± 70 bp s^−1^ (mean ± SEM) with a distribution that reflects intrinsic differences among individual replisomes (Figure 2E). These rates are similar to those obtained before in ensemble (45) and single-molecule experiments (33,37,43,46).

**Figure 2.**
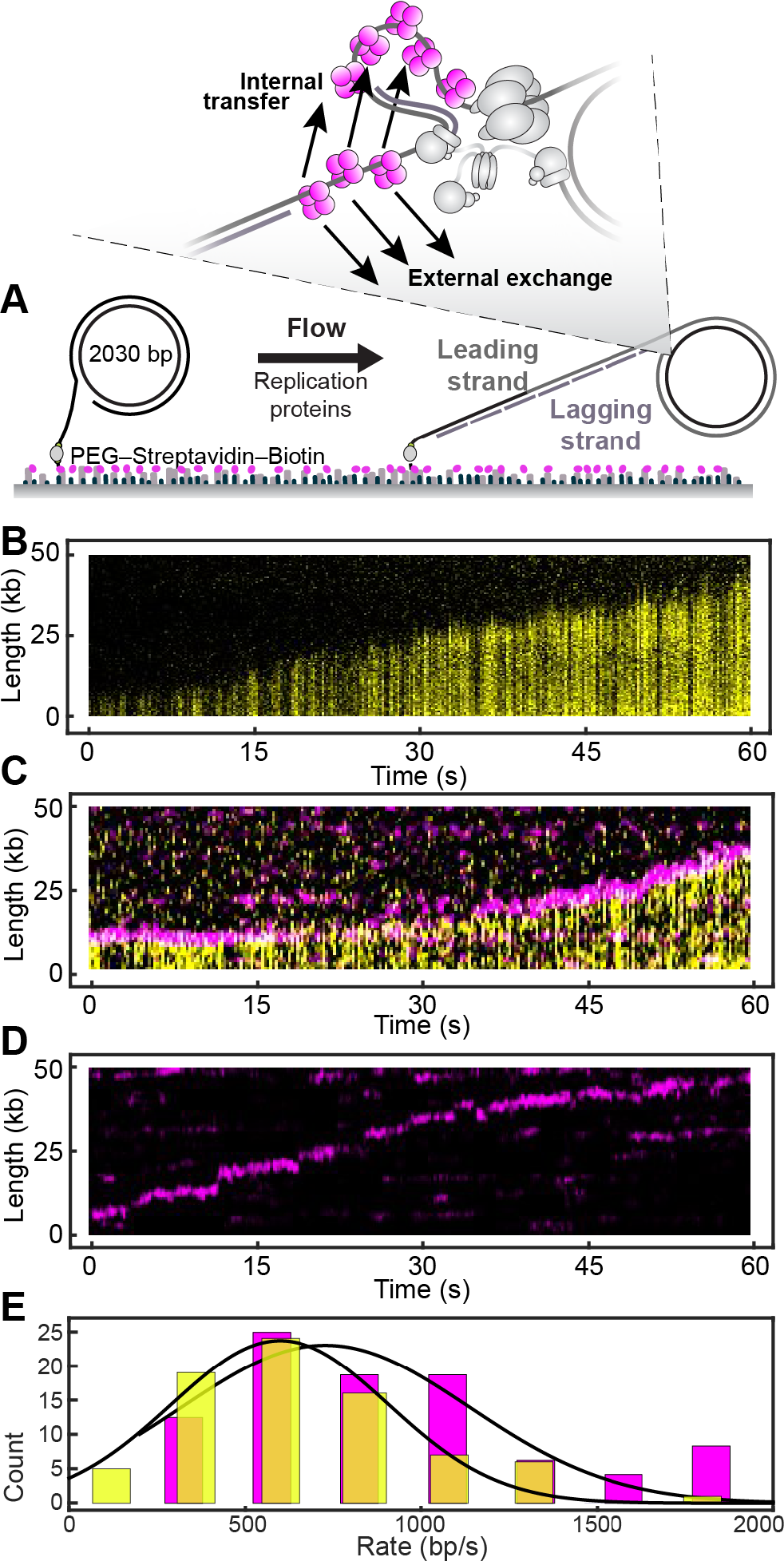
Visualisation of SSB in the single-molecule rolling-circle assay. (**A**) Schematic representation of the experimental design. 5′-Biotinylated circular DNA is coupled to the passivated surface of a microfluidic flow cell through a streptavidin linkage. Addition of the *E. coli* replication proteins and nucleotides initiates DNA synthesis. The DNA products are elongated hydrodynamically by flow, labelled with intercalating DNA stain, and visualised using fluorescence microscopy while replication takes place (37). The schematics are not to scale, with the circles resulting in a diffraction-limited spot in the imaging and the product DNA showing up as long lines. (**B**) Kymograph of an individual DNA molecule undergoing coupled leading-and lagging-strand replication. The yellow indicates the fluorescence intensity of stained DNA. (**C**) Representative kymograph of simultaneous staining of double-stranded DNA and fluorescence imaging of labelled SSB (red) in real time. The kymograph demonstrates that the fluorescent spot corresponding to SSB co-localises with the tip of the growing DNA product, at the position of the circle and its replication fork. (**D**) Kymograph of the red-labelled SSBs on an individual DNA molecule. The intensity of the SSB signal does not photobleach but instead remains constant for the duration of the experiment, indicating at least some SSB is exchanged. (**E**) Histograms of the rate of replication for wild-type SSB (626 ± 73 bp s^−1^, *N* = 71) and labelled SSB (720 ± 55 bp s^−1^, *N* = 59) fit to Gaussian distributions. The similarity between these rates shows that the label does not affect the behaviour of SSB during replication. See also Supplementary Figure S2.

To visualise the behaviour of SSB during rolling-circle replication, we labelled a mutant of SSB containing a single cysteine (SSB-K43C) with a red fluorophore (AlexaFluor 647; Supplementary Figure S2A). The labelled SSB was active in leading-and lagging-strand synthesis, producing Okazaki fragments of size distributions identical to those obtained with wild-type SSB in an ensemble-averaging solution-phase reaction (Supplementary Figure S2C). We then used the fluorescently labelled SSB at a concentration of 20 nM (all SSB concentrations are as tetramers) in the rolling-circle assay. Simultaneous imaging of the stained DNA and labelled SSB showed that the SSB is located at the tip of the growing DNA, consistent with the labelled protein being integrated into active, reconstituted replisomes (Figure 2C). The single-molecule replication rates in the presence of the labelled SSB were similar to those using unlabelled wild-type SSB (Figure 2D and E), in agreement with our ensemble assays. Thus, the label does not affect the behaviour of SSB in a fully reconstituted DNA-replication reaction supporting simultaneous leading-and lagging-strand synthesis. We have reported previously that under the same conditions, polymerases bind to gaps between Okazaki fragments behind the replication fork (33). Interestingly, we do not observe SSB signals at stationary positions on the product DNA, behaviour that would be expected to result in horizontal lines in the kymographs as observed before for labelled Pol III* retained at the ends of completed Okazaki fragments in the absence of their further processing (33). There has been debate over many years about whether lagging-strand polymerases may be recycled to new primer termini before (signalling model) or on Okazaki fragment completion (collision model), most recently summarised by Benkovic and Spiering (47). Our observation of the absence of SSB in gaps does not discriminate between these models because we are unable to resolve SSB left transiently in the wake of the replisome from SSB at the fork; the two populations would have to be separated well beyond the diffraction limit, or by at least 500 nm (corresponding to ~1,800 bp). During the time it takes the replisome to cover this distance, polymerases recruited from solution will have filled gaps in any incomplete Okazaki fragments (33). The absence of SSB spots behind the replisome does suggest that completion of an Okazaki fragment does not leave a ssDNA gap sufficiently large (>35 nt) for SSB to bind.

The intensity of the fluorescence signal from the SSB at the replisome remains essentially constant throughout the experiment (Figure 2D). If all SSB molecules were internally recycled and retained in the replisome, the fluorescence intensity should decay at the characteristic lifetime of photobleaching, which is 9.5 ± 0.8 s under these conditions (Supplementary Figure S2B). Therefore, at least some SSB molecules at the fork are replaced by new ones from solution. This exchange needs to take place at a rate that is high enough to keep the steady-state level of unbleached SSB sufficient to be observable.

### Dynamic behaviour of SSB *in vitro*

We next used *in vitro* single-molecule FRAP (fluorescence recovery after photobleaching) experiments (33) to quantify the dynamic behaviour of SSB during DNA replication, using the same rolling-circle reaction (Figure 2A). Instead of continuous imaging at constant laser power, we periodically bleached all SSB in the field of view using 100-fold higher power (Figure 3A, left). Due to the buffer flow and high diffusional mobility, bleached SSB that is free in solution will rapidly move away and be replaced by unbleached, bright SSB. After the photobleaching pulse, we monitor the recovery of the fluorescence signal at the replisome as a readout for the kinetics of introduction of new, unbleached SSB at the replication fork. This measurement allows us to distinguish between internal transfer and external exchange of SSB: If SSB were transferred internally and retained at the fork, the fluorescence would not recover after photobleaching (Figure 3A, top right); if dark, bleached SSB is exchanged with fluorescent SSB from solution, however, we should observe recovery of the fluorescence intensity at the fork. Figure 3B shows a kymograph of a FRAP experiment using 10 nM labelled SSB. The vertical lines correspond to the high-intensity FRAP pulses. After each pulse, the fluorescence of the SSB spot decreases to zero as the population is bleached. This bleaching is followed by a gradual increase in intensity, indicating that SSB from solution associates at the fork. We determined the intensity after each FRAP pulse over time by averaging over *N* = 24 replisomes (Figure 3C). At 10 nM SSB, we find the recovery time (τ) to be 10 ± 1 s (Figure 3D). We then repeated this measurement for SSB concentrations varying from 2 to 100 nM (Figure 3D and E). At 2 nM SSB, the fluorescence signal recovers slowly (τ = 20 ± 7 s, *N* = 20), while at 100 nM, its recovery is ~10-fold faster (τ = 2.9 ± 1.7 s, *N* = 18). These data show that SSB exchange is concentration dependent, with faster exchange occurring at higher concentrations.

**Figure 3.**
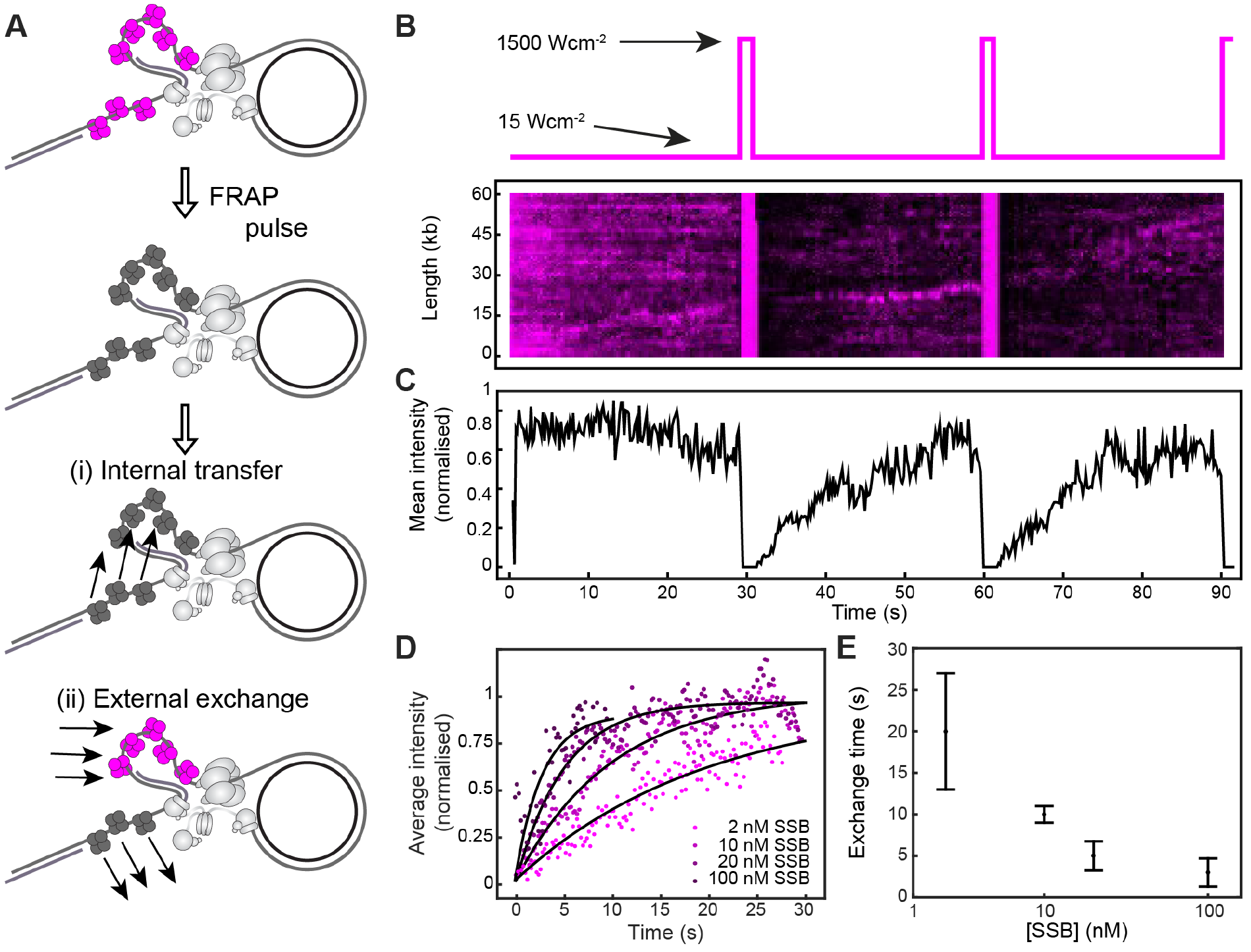
Quantification of the SSB exchange time using single-molecule FRAP. (**A**) Schematic representation of the FRAP experiments. SSB molecules are initially in a bright state (top). After a high intensity FRAP pulse, all SSB in the field of view is photobleached. If SSB is internally transferred, no fluorescence recovery should be observed (i). If SSB is externally exchanged, the fluorescence should recover rapidly (ii). (**B**) Imaging sequence used during the FRAP experiments (top panel). A representative kymograph of labelled SSB at the replication fork (bottom panel) in a FRAP experiment. After each FRAP pulse (indicated by the vertical red line) all SSB molecules have bleached. The fluorescence intensity recovers as unbleached SSB exchanges into the replisome. (**C**) The average intensity over time from 20 replisomes with 10 nM SSB in solution. (**D**) The first three recovery phases in (C) were averaged again to give the final averaged normalised intensity over time after a FRAP pulse. This curve was then fit to provide a characteristic exchange time. This procedure was repeated for four concentrations of SSB ranging from 2-100 nM. (**E**) Exchange time as a function of SSB concentration shows a concentration-dependent exchange time. See also Supplementary Figure S3.

### SSB is recycled for multiple Okazaki fragments

Having obtained information on the time scale of SSB turnover at the replisome, we then characterised the number of Okazaki fragment priming and synthesis cycles that occur during that time window. We did so by determining rates of replication and the lengths of the Okazaki fragments. First, we used the single-molecule rolling-circle assay to obtain the DNA replication rates at the different SSB concentrations we used in the FRAP experiments. At all concentrations of SSB, the replication rate was around 750 bp s^−1^, with no statistically significant differences (Supplementary Figure S3). The observation that SSB recovery times can be as high as tens of seconds (Figure 3E) suggests that the protein is recycled within the replisome for a period that corresponds to the synthesis of many thousands of base pairs. With an Okazaki-fragment length of 1–2 kb (48), our observations suggest that the replisome retains the SSB for a duration well beyond the time need to synthesise an Okazaki fragment. Such a long retention time can only be explained by a mechanism that allows internal transfer of SSB from one Okazaki fragment to the next.

To verify this interpretation, we measured the length of Okazaki fragments generated under our conditions, using both an ensemble-averaging biochemical approach and direct single-molecule observation. It has previously been reported that the SSB concentration has an effect on Okazaki-fragment length (49). To recapitulate this concentration effect, we first performed ensemble rolling-circle replication experiments. Replication reactions containing all proteins required to support simultaneous leading-and lagging-strand synthesis, with SSB at different concentrations, were allowed to proceed for 30 min. The resulting products were separated on an alkaline agarose gel and stained with an ssDNA stain for visualisation (Figure 4A). The intensity distributions were normalised to correct for the fact that the intrinsic intensity per mole of product DNA scales linearly with length. The product length distributions show that the Okazaki fragments are shorter for lower SSB concentrations (1.4 ± 0.2 knt at 2 nM versus 2.8 ± 1.0 knt at 200 nM), a two-fold difference in product length for a 100-fold change in SSB concentration.

**Figure 4.**
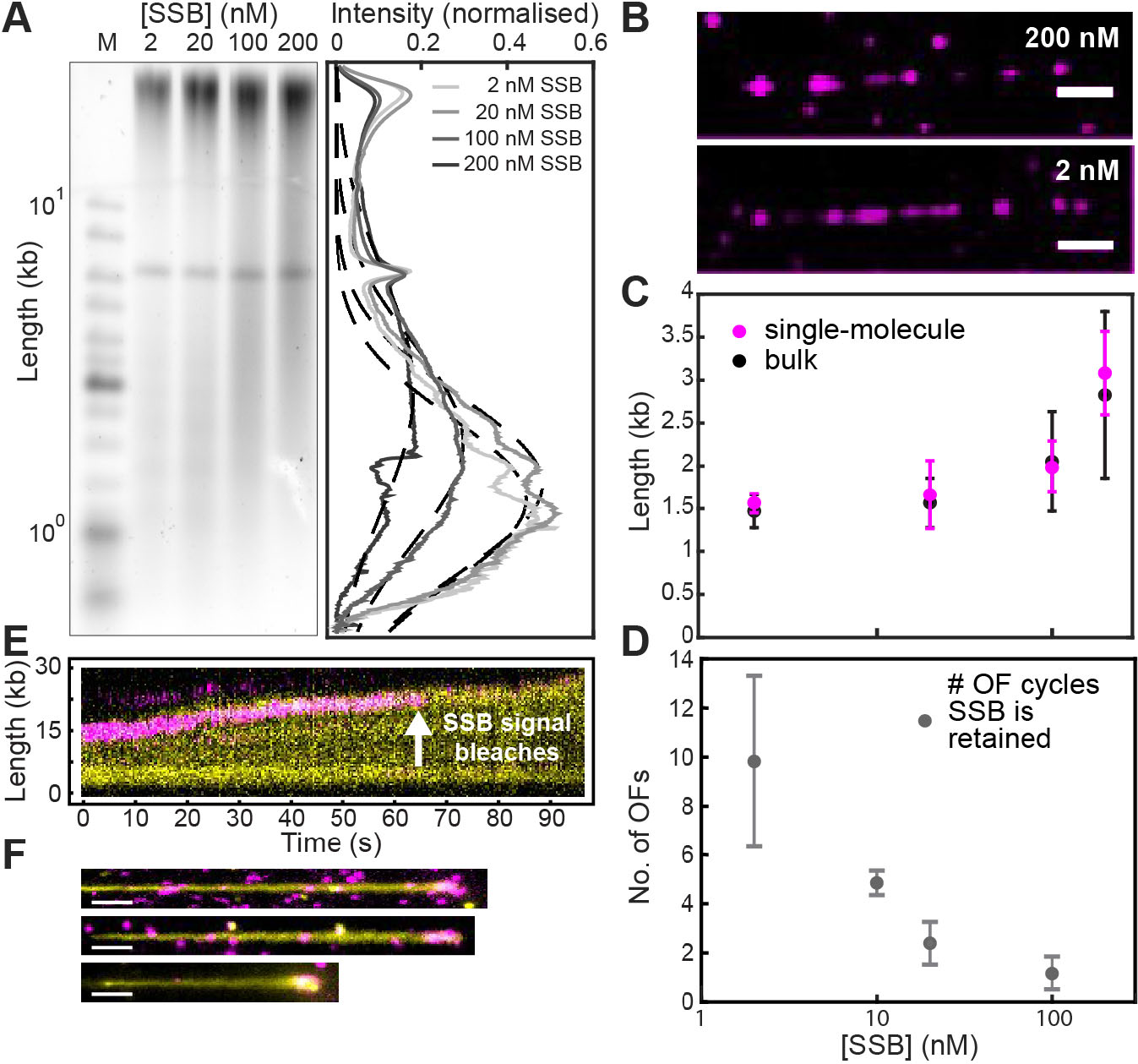
Internal transfer of SSB. (**A**) Alkaline agarose gel of M13 rolling circles replicated using concentrations of SSB identical to those used in the FRAP experiments (left panel). The right panel shows the intensity profiles of lanes 2-5. The Okazaki fragment size distributions are centred at 1.4 ± 0.2, 1.5 ± 0.3, 2.0 ± 0.6 and 2.8 ± 1.0 kb (mean ± standard deviation), in the presence of 2, 10, 20 and 100 nM SSB, respectively. Intensity profiles have been corrected for the intrinsic difference in intensity of different size fragments using the ladder as a size standard. (**B**) Representative images showing SSB bound in the gaps between Okazaki fragments. Replication was carried out using a polymerase pre-assembly assay, with different concentrations of labelled SSB (top: 200 nM; bottom: 2 nM). Since there is no polymerase in solution to fill in the gaps between nascent Okazaki fragments, SSB will bind there. Therefore, the distance between two SSB spots is a measure for Okazaki-fragment length. All unbound proteins were washed out for imaging. The scale bar represents 10 kb. (**C**) Comparison of Okazaki-fragment lengths measured in the ensemble assay described in panel (A) (black) and at the single-molecule level (red, Supplementary Figure S4). (**D**) The number of Okazaki-fragment (OF) synthesis cycles that are supported by the same pool of SSB, as a function of SSB concentration. The numbers were obtained by dividing the SSB recovery times in Figure 3E by the time it takes to synthesise one Okazaki fragment using the lengths found in (C). (**E**) Kymograph of the simultaneous imaging of DNA and pre-assembled SSB, in the absence of SSB in solution. Replication was initiated in the presence of SSB and synthesis was allowed to proceed for 1 min. Then, indicated by *t* = 0, the solution was exchanged to a buffer containing all replication components, but omitting SSB. Leading-and lagging-strand synthesis continues after the SSB signal photobleaches, suggesting that SSB is still present at the fork. (**F**) Representative examples of long DNA products with labelled SSB present at the fork upon conclusion of an SSB pre-assembly experiment (scale bar is 10 kb). See also Supplementary Figures S4 and S5.

It can be argued that the effect of SSB concentration on the Okazaki-fragment length may be different when using the single-molecule assay. In these experiments, SSB is continuously replenished through the buffer flow, whereas in an ensemble experiment SSB would be sequestered from solution if more ssDNA were generated. To test whether such a difference exists, we measured the Okazaki-fragment lengths in our single-molecule assays. However, in our continuous-flow rolling-circle experiments using DNA staining, we do not have the spatial resolution to observe the gaps between Okazaki fragments in the product DNA. Furthermore, as discussed above, DNA polymerases in solution rapidly fill in the gaps between the Okazaki fragments, making them too small for SSB to bind, thus preventing us from using fluorescent SSB to detect junctions between Okazaki fragments. To resolve this issue, we use conditions that prevent free polymerases from filling in Okazaki-fragment gaps. This is achieved by use of pre-assembly experiments, in which polymerases are present in solution during the initial phase of establishing replisomes at the forks, but left out during the phase in which the pre-assembled replication complexes produce DNA. Such a design forces the replisome to retain the polymerase holoenzyme (33) and allows labelled SSB to bind ssDNA gaps between the Okazaki fragments without being displaced by other DNA polymerases (Figure 4B). The distance between the SSB spots can then be used as a measure of Okazaki-fragment length. By measuring the distances between many pairs of SSB spots, we obtained distributions of distances for different SSB concentrations, which were fit with single-exponential decay functions to obtain the average Okazaki-fragment lengths (Supplementary Figure S4A). These lengths are the same as those measured in the ensemble experiment, showing that the Okazaki-fragment distributions are similar between the two experiments, with a similar dependence on SSB concentration (Figure 4C).

We can now use the single-molecule observations of Okazaki-fragment length for different SSB concentrations to directly compare the time required for the replisome to synthesise a single fragment to the time associated with SSB recovery. Converting the information on Okazaki-fragment lengths (Figure 4) into times by using the replication rate (Supplementary Figure S3), and by dividing SSB recovery times (Figure 3E) by this Okazaki-fragment time, we determine the number of Okazaki-fragment synthesis cycles that are supported by the same pool of SSB (Figure 4D). This analysis shows that at low concentrations, SSB is retained within the replisome during synthesis of multiple (~10) Okazaki fragments. This number decreases as the SSB concentration is increased, further suggesting a concentration-dependent competition between internal transfer and external exchange.

As SSB is continuously displaced from the ssDNA by the lagging-strand polymerase, retention must mean that SSB molecules are transferred internally to newly-exposed ssDNA behind the helicase. To see if we could push the equilibrium between internal transfer and external exchange completely towards internal transfer, we carried out a pre-assembly experiment eliminating all free SSB from solution. In this assay, replication was initiated in the presence of labelled SSB and allowed to proceed for 1 min. We then switched to a buffer containing all replication proteins, but omitting SSB and thereby preventing any external exchange. Simultaneous imaging of the stained product DNA and the labelled SSB shows that the DNA tail keeps growing after the SSB signal disappears due to photobleaching of the dye (Figure 4E). We conclude that under these conditions, the lifetime of SSB on ssDNA is much longer than the photobleaching lifetime of around 10 s (Supplementary Figure S2B). In support of this observation, we next imaged the long DNA product molecules only after the replication reaction had finished rather than illuminating continuously, and observed labelled SSB foci were still present at the tip (Figure 4F). To calculate the number of SSB tetramers present on the ends of these product molecules, we measured the intensity of these spots. When we divide their average intensity by the intensity of a single tetramer (Supplementary Figure S4B), we find that the average number of SSBs stably bound at the end of the DNA products corresponds to 35 ± 3 tetramers (Supplementary Figure S4C). This number of SSB tetramers corresponds to a ssDNA footprint of slightly more than 1 kb (assuming 35 nt per SSB tetramer), the same length scale as an Okazaki fragment. Remarkably, this observation suggests that upon removal of SSB from solution, the replisome retains its original complement of SSB for many tens of kb of synthesis, supporting that internal transfer can be highly efficient in the absence of SSB in solution.

As a final control experiment to test if internal transfer may be even more efficient with wild-type SSB than with the labelled SSB-K43C mutant protein, we allowed initial replisome assembly and 1 min of synthesis in the presence of 10 nM unlabelled wild-type SSB. We then switched to 10 nM labelled SSB in the buffer flow and monitored the increase in fluorescence intensity at the fork, as labelled SSBs exchange into the replisome (Supplementary Figure S5). The labelled SSBs exchanged into the replisome on a time scale that was similar to that measured in Figure 3E. This observation indicates that the internal-transfer efficiencies of wild-type and labelled SSB-K43C SSBs are similar.

### Dynamic behaviour of SSB *in vivo*

To extend our *in vitro* observations, we used *in vivo* single-molecule FRAP experiments to study the dynamics of SSB in live *E. coli* cells. *In vivo* FRAP has previously been used to measure the dynamics of other replisome components including the Pol III holoenzyme and the DnaB helicase (50). We used *E. coli* cells in which the chromosomal *ssb* gene is supplemented by a gene that generates a C-terminal fusion of the protein with a yellow fluorescent protein (YPet) (51). We verified by PCR that both the wild-type *ssb* gene and the *ssb-YPet* fusion gene are present in this strain (Supplementary Figure S1B). On expression in this strain, it is assumed that SSB and SSB-YPet form mixed tetramers. To confirm that the mixed tetramers are functional, we first showed that the growth rate of the *ssb-YPet* cells is similar to that of isogenic wild-type *E. coli* (Supplementary Figure S1A). Next, to confirm that the labelled SSB forms part of active replisomes, we studied co-localisation of SSB-YPet and DnaQ-mKate2, producing red-labelled Pol III, in a dual-colour strain expressing both. We found that as many as 100% and on average 67% of DnaQ foci per cell co-localise with SSB foci (*N* = 65 cells).

Measurement of the fluorescence recovery of SSB within cellular foci requires the ability to specifically bleach the fluorescence within a single replisome focus without bleaching the SSB in the rest of the cell. To this end, we placed a pinhole in a motorised filter wheel in the excitation path, producing a tight, diffraction-limited excitation focus (full width at half maximum, 500 nm). Using this pinhole and a high laser power (200 W cm^−2^), we can bleach a single focus with high spatial specificity (Figure 5A). The subsequent fluorescence recovery was visualised by lowering the laser power (to 2 W cm^−2^) and by moving the pinhole out of the beam path. Figure 5B shows bleaching and recovery of a SSB-YPet focus within a single cell (green arrow). The first frame was acquired before applying the FRAP pulse. The image acquired immediately after the pulse (*t* = 0 s) shows that the fluorescence from the single focus has bleached, while the SSB in the cytosol remains fluorescent. In subsequent frames, we see that the fluorescence recovers, indicating that non-bleached SSB from the cytosol exchange into the focus. To quantify the exchange time, we measure the intensity of the foci over time after the bleaching pulse. An average intensity trajectory (*N* = 29 foci) shows an initial recovery of the fluorescence after the photobleaching pulse, followed by a decay in intensity (Figure 5C) due to photobleaching of the YPet probe during visualisation, even at the lower imaging intensities after the high-intensity bleaching pulse. To correct for this, we measured the average photobleaching behaviour of the probe by monitoring the fluorescence from other cells within the same field of view (Figure 5D). Since these cells were not subject to the high-power bleaching pulse, their fluorescence signals provide an internal benchmark for the gradual photobleaching induced by the lower-power imaging illumination. These photobleaching data were fit with a single-exponential decay function (green line). This fit was then used to correct the FRAP intensity trajectories, with the corrected trajectory showing behaviour that is now representative of the recovery of the pool of unbleached SSB at the replisomal spot (Figure 5E). By fitting these recovery data (green line), we obtain a recovery time of 2.5 ± 1.7 s. This value is similar to the time scale we obtained from the *in vitro* experiments at high SSB concentrations, a similarity that was expected since the estimated concentration of SSB *in vivo* is 300–600 nM during mid-log growth. Assuming that Okazaki fragments produced in the cell are 1–2 knt in length (52) and the replication rate is ~1000 bp/s, such an exchange time would suggest that for every Okazaki fragment cycle, roughly half of the SSB is internally recycled for the next fragment and the other half is exchanged with free SSB.

**Figure 5.**
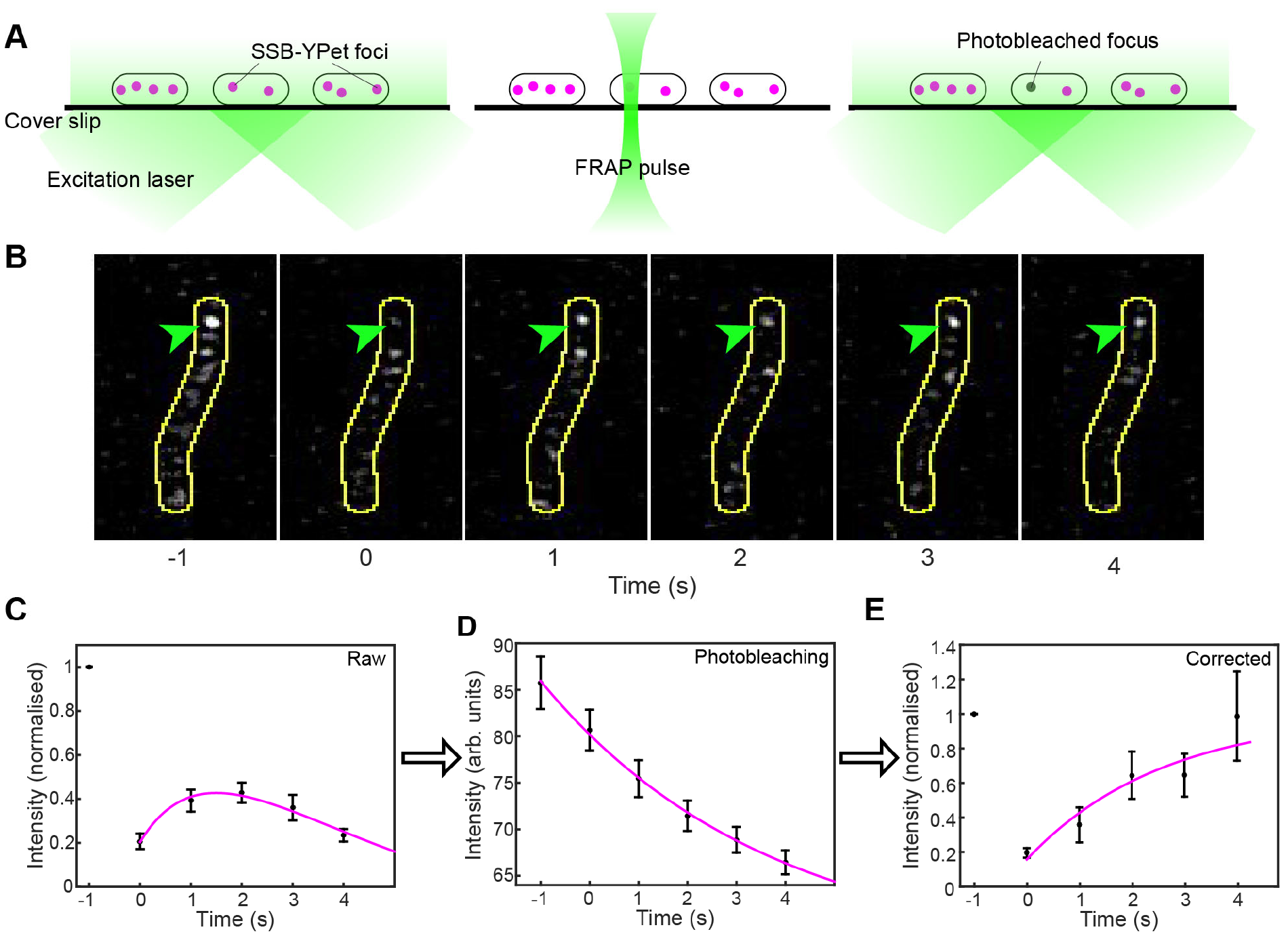
Visualisation of SSB dynamics *in vivo*. (**A**) Schematic representation of the *in vivo* FRAP setup. SSB-YPet foci (red) are visualised before FRAP (left). By placing a pinhole in the beam path, a single focus will be darkened without bleaching cytosolic SSB-YPet (middle). After the FRAP pulse the recovery of fluorescence can be monitored. (**B**) Representative images of *in vivo* FRAP experiments. At times before *t* = 0, the focus (indicated by the green arrow) is bleached using a high-intensity FRAP pulse. The fluorescence recovers as fluorescent SSB from the cytosol exchanges into the replisome focus. The cell boundaries are indicated by the yellow line. (**C**) Averaged normalised FRAP intensity trajectory (*N* = 29). After initial recovery, the fluorescence intensity decreases due to photobleaching. (**D**) Average intensity over time for *ssb-YPet* cells outside of the FRAP volume (*N* = 40). These data were fit with a single-exponential decay function (green line) to obtain the photobleaching lifetime. (**E**) Averaged normalised FRAP intensity trajectory, corrected for photobleaching. The green line represents a fit to the data, from which we obtained the characteristic *in vivo* exchange time for SSB (τ = 2.5 ± 1.7 s). See also Supplementary Figure S1.

## DISCUSSION

Biochemical studies suggest two different mechanisms to describe how SSB binds to and dissociates from ssDNA within the replisome. In the external-exchange model, newly exposed ssDNA is bound by SSB from solution. In an alternative model, SSB is recycled within the replisome through an internal-transfer mechanism. Using an *in vitro* single-molecule visualisation approach, we show here that SSB can be recycled within the replisome on time scales corresponding to the synthesis of multiple Okazaki fragments, thereby verifying the existence of an internal-transfer mechanism of SSB molecules at the fork. At higher SSB concentrations, however, we observe that this mechanism is in competition with external exchange of SSB with molecules present in solution. Using single-molecule imaging of labelled SSB in live bacterial cells, we show that both processes occur at the replication fork in a cellular context and that roughly half of the SSB is internally recycled for the next Okazaki fragment.

We conclude that the *E. coli* replisome strikes a balance between internal transfer and external exchange of SSB. In the absence of SSB in solution, the original population is retained within the replisome and is efficiently recycled from one Okazaki fragment to the next (Figure 4E). The existence of such an internal-transfer mechanism has been hypothesised, as it has been shown that SSBs can be transferred between DNA strands through a transient paired intermediate (13,14,24). Internal transfer has, however, not been shown before in the context of active DNA replication. We show here that in the absence of competing free SSB in solution, SSB is recycled by the replisome for many tens of kb. Estimates of the total concentration of SSB tetramers in *E. coli* cells have ranged from 50-600 nM (300–600 nM during mid-log phase), which would be sufficient to coat up to ~20 knt of ssDNA in the SSB_35_ binding mode (2,13,21). The concentration of available SSB within the cytosol could be significantly lower with SSB bound to the various ssDNA substrates within the cell. Moreover, at high growth rates, the cell could contain up to 12 active replisomes (25), leaving little free SSB. This lack of readily available SSB may make its binding from solution too slow to coat the rapidly produced ssDNA at replication forks, resulting in exposure of vulnerable ssDNA that can be nucleolytically attacked, can form secondary structures, or can act as a substrate for ssDNA-binding proteins that trigger undesired pathways or responses (*e.g.*, RecA). The internal-transfer mechanism could be a way to ensure rapid SSB coating of newly exposed ssDNA, thereby allowing replication to continue at normal rates without creating large amounts of naked ssDNA.

While it is not known what proportion of SSB *in vivo* is in either of the predominant binding modes (Figure 1B), internal transfer without equilibration with SSB in solution presumably requires it to be in the SSB_35_ mode. We propose that the advancing lagging-strand polymerase is capable if necessary of converting SSB from the SSB_65_ to the SSB_35_ mode by reducing the availability of ssDNA, enabling its co-operative transfer to ssDNA generated behind the DnaB helicase.

In the presence of competing SSB in solution, however, this internal-transfer mechanism is in competition with external exchange of SSB at a rate that is dependent on its concentration in solution (Figure 3E). Such a concentration-dependent exchange mechanism has recently been observed for other proteins that form part of multi-protein complexes (33,44,53–60). Under highly diluted conditions, these proteins can remain stably bound within the complex for long periods of time. Yet, rapid (sub-second) exchange is observed at nanomolar concentrations. Such concentration-dependent dissociation can be explained (61) and mathematically described (62–65) by a multi-site exchange mechanism in which a protein is associated with a complex *via* multiple weak binding sites, as opposed to a single strong one.

A competition between stability and plasticity that depends on concentration seems harder to comprehend for SSB. Under any circumstance, dilute or not, the SSB–ssDNA interaction has to be disrupted as new dsDNA is synthesised on the lagging strand. Therefore, stability, defined as retention within the replisome, cannot be achieved in the same way as described for other proteins, but instead needs to rely on a mechanism of internal transfer. The disruption of the SSB–DNA interaction due to lagging-strand synthesis would be followed by rapid rebinding of SSB to the next Okazaki-fragment template produced behind the helicase, thereby preventing dissociation of the SSB from the replisome. If, however, there are competing SSB molecules in close proximity to the fork, one of these can bind at the newly exposed ssDNA, thereby blocking that binding site. Consequently, SSB molecules released from the lagging strand can no longer rebind and are effectively competed out of the replisome.

Our observations of SSB dynamics in living cells are consistent with the hypothesis that both internal transfer and external exchange are physiologically relevant pathways accessible to the replisome during coupled DNA replication. In our measurements, during mid-log growth (estimated intracellular SSB concentrations of 300–600 nM), the balance seems to be towards external exchange, with relatively fast exchange times of 2.5 ± 1.7 s. This time scale is consistent with those we obtained in our *in vitro* measurements.

A multi-site exchange mechanism confers both stability and plasticity to the replication machinery, allowing the replisome to operate under different cellular conditions. Our work, combined with other recently published studies, presents a much more dynamic picture of the replisome, distinctly different from the deterministic models generated over the last few decades. It is important to point out that the stochasticity and plasticity observed in recent single-molecule experiments are all consistent with fundamental chemical principles and can be readily explained by hierarchies of weak and strong intra-replisomal interactions (66). The apparent generality of the models emerging from these studies suggests that the behaviours of other complex multi-protein systems might also be governed by such exchange processes and might suggest that evolution of complex interaction networks has arrived at an optimal balance between stability and plasticity.

## Supporting information

## SOFTWARE AVAILABILITY

Home-built ImageJ plugins have been deposited on the Github repository for Single-molecule/Image analysis tools (https://github.com/SingleMolecule).

## SUPPLEMENTARY DATA

Supplementary Data are available at NAR online.

## ACKNOWLEDGEMENT

The authors thank Sarah Henrikus, Megan Cherry and Enrico Monachino for providing protocols and reagents and Bénédicte Michel for kindly providing the *ssb-YPet* strain.

