## Supplementary material for "Recycling of single-stranded DNA-binding protein by the bacterial replisome"

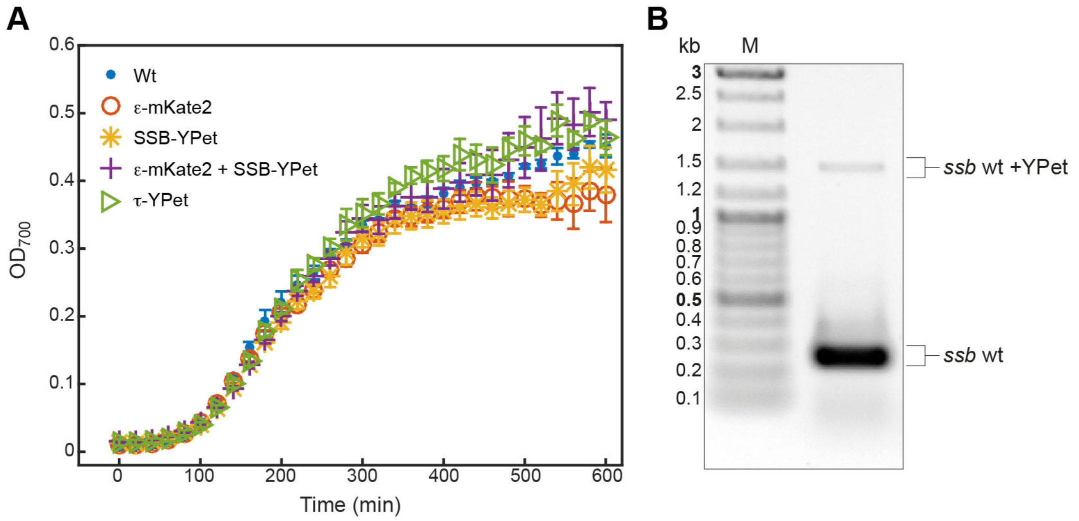


**Supplementary Figure S1.** *ssb–YPet* strain JJC5380. (**A**) Growth curves for *E. coli* strains: wild-type *E. coli* (blue), cells expressing C-terminal derivatives of ε (*dnaQ-mKate2*, red), SSB (*ssb-YPet*, JJC5380, yellow), both ε and SSB (*dnaQ-mKate2* + *ssb-YPet*, purple), and τ (*dnaX-YPet*, green) subunits under control of their endogenous promoters. Growth curves were measured for 10 h. Experiments were performed in triplicate. The errors represent the experimental variation. (**B**) The presence of both the fusion and wild-type *ssb* genes was checked by PCR using oligonucleotides 5'-GCAGGGTGGCAATCAGTTCAG-3' and 5'-GTTTCCGCCTGTTGGTTCGCA-3' and chromosomal DNA as template. The gel of the PCR products shows both the wild-type *ssb* and the *ssb–YPet* fusion are present.

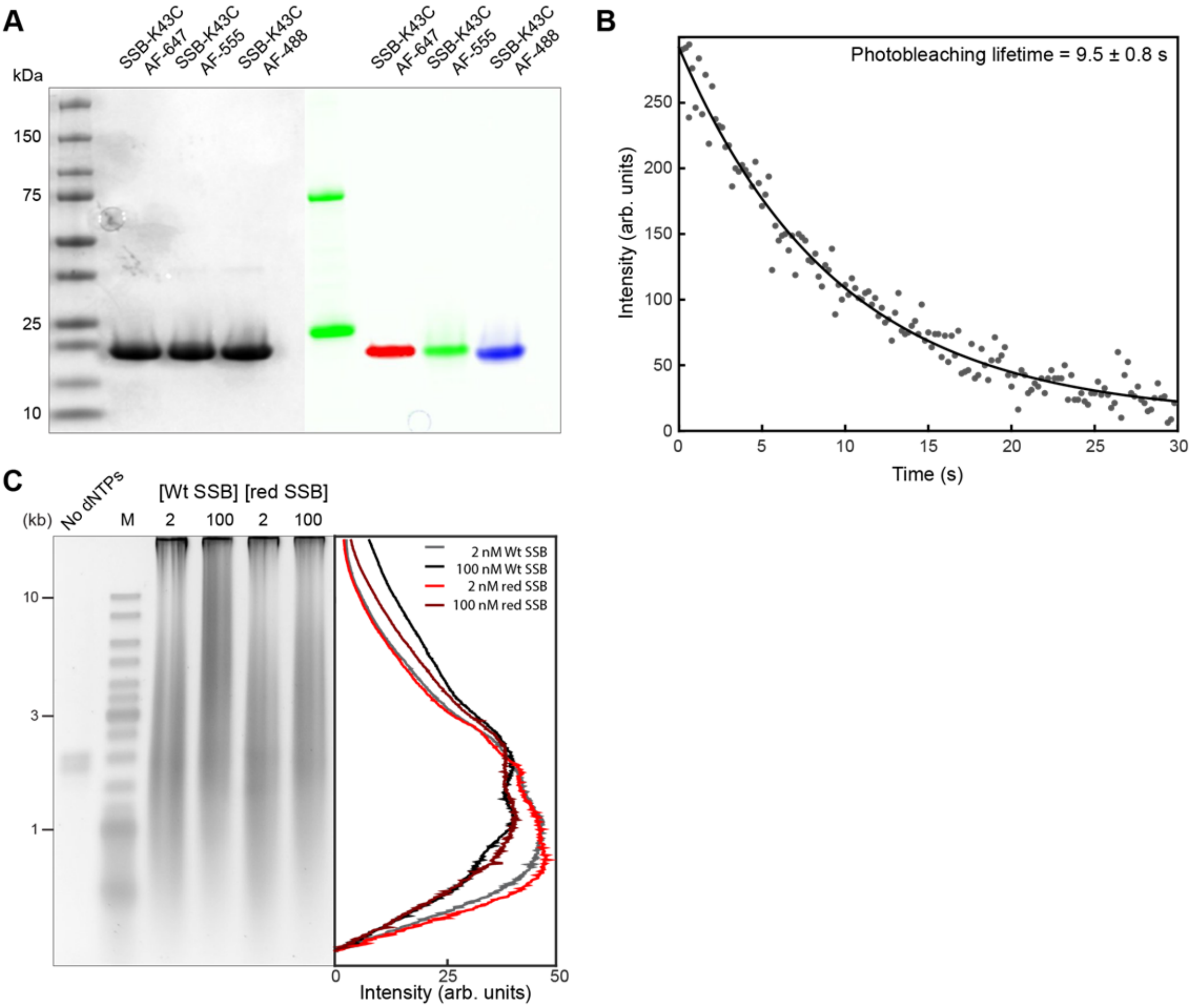


**Supplementary Figure S2.** Fluorescently-labelled SSB. (**A**) SDS-PAGE of labelled SSB-K43C. Lanes on the left are stained with Coomassie blue, while those on the right are unstained. (**B**) Average photo-bleaching trajectory for SSB-AF647 ($N$ = 4 fields of view, 568 molecules) at excitation power density of 700 mW cm$^{-2}$. From a fit with single-exponential decay function (black line), we obtained a photobleaching lifetime of 9.5 ± 0.8 s. (**C**) Comparison of activities of wild-type and labelled SSBs. (left) Alkaline agarose gel of coupled DNA replication. Reactions were performed on a 2-kb circular dsDNA template with 2 and 100 nM of either wild-type (wt) SSB or red-labelled SSB. The gel was stained with SYBR-Gold. (right) Intensity profiles of lanes 3–6. Intensity profiles have been corrected for the difference in intensity of different sized fragments using the ladder as a standard.

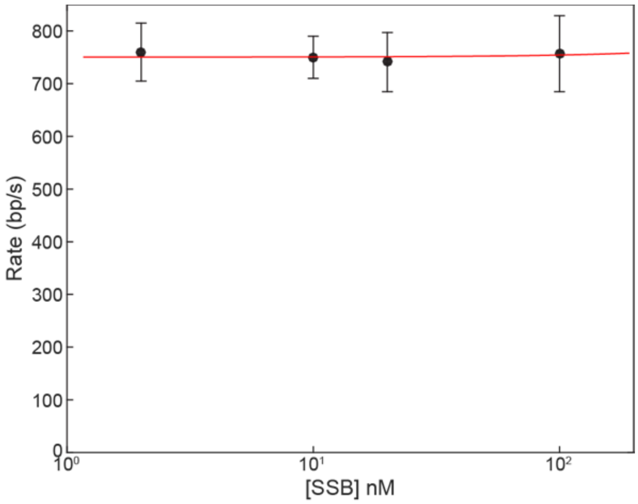


**Supplementary Figure S3.** Rate of replication is independent of SSB concentration. Replication rate distributions were obtained and fitted as described in Figure 2. The points represent the mean of the distribution, and the error bars are the SEM. The red line represents a linear fit to the data giving a rate of 750 ± 14 bp $s^{-1}$ (error is error of the fit).

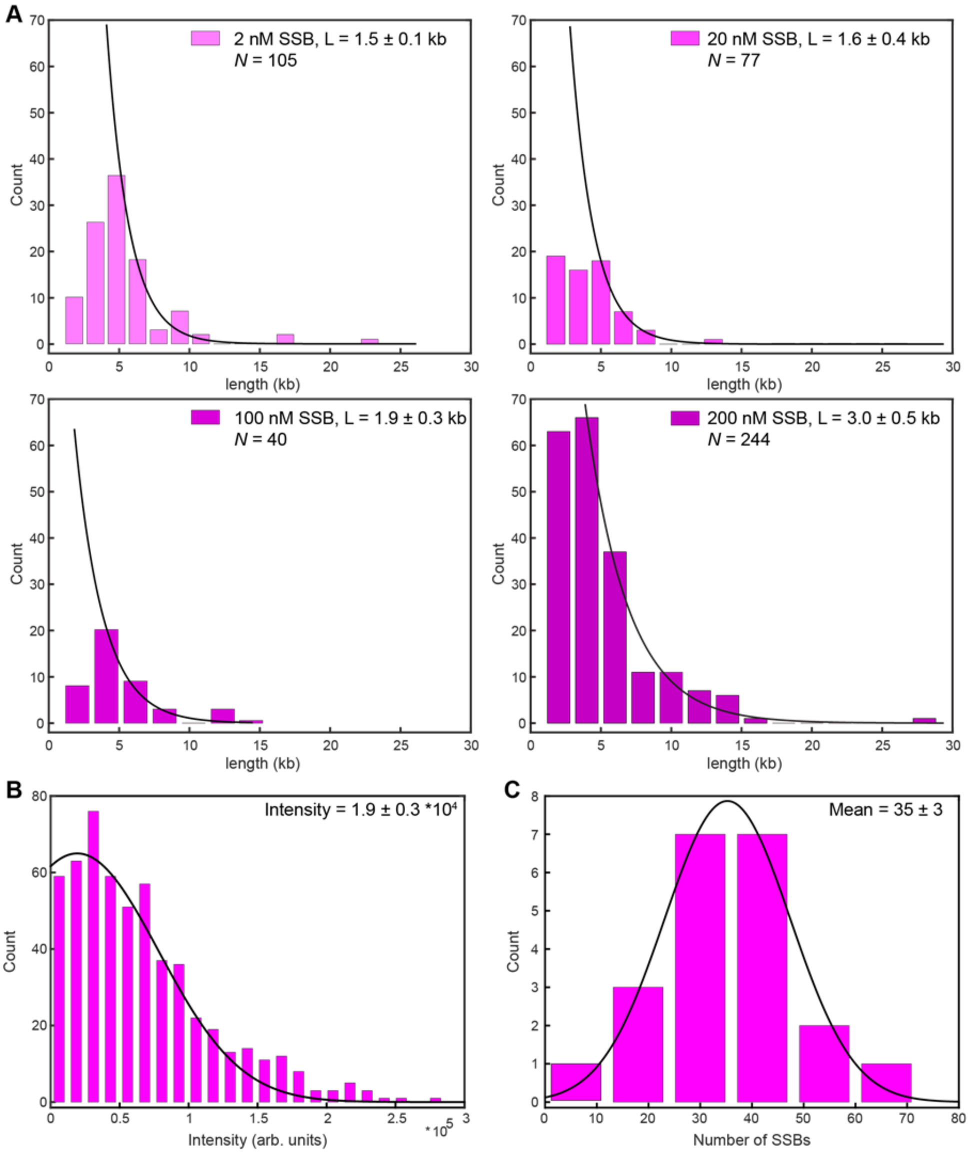


**Supplementary Figure S4.** Okazaki fragment sizes and quantification of SSB. (**A**) Single-molecule measurement of Okazaki-fragment length for different concentrations of SSB. The histograms represent distributions of distances measured between SSB spots. The black lines are a single-exponential fit to the data. The first bars are not included in the fit to take into account under-sampling at distances shorter than or comparable to the diffraction limit. (**B**) Histogram of the intensity distribution of single SSB molecules. The average intensity of a single labelled SSB was calculated by immobilisation on the surface of a cleaned microscope coverslip in imaging buffer. The imaging was under the same conditions as used during the single-molecule rolling-circle experiments. Using ImageJ with in-house built plugins, we calculated the integrated intensity for every SSB in a field of view after applying a local background subtraction. The histogram was fit with a Gaussian distribution function to give a mean intensity of $(1.9 \pm 0.3) \cdot 10^4$. The error represents the standard error of the mean. (**C**) Histogram of the number of SSBs at the fork at the conclusion of an SSB pre-assembly experiment. The numbers were obtained by dividing the intensities at the fork by the intensity of a single SSB found in (B). From the Gaussian fit (black line), we find that there are 35 ± 3 (mean ± SEM) SSB molecules at the fork (*N* = 31).

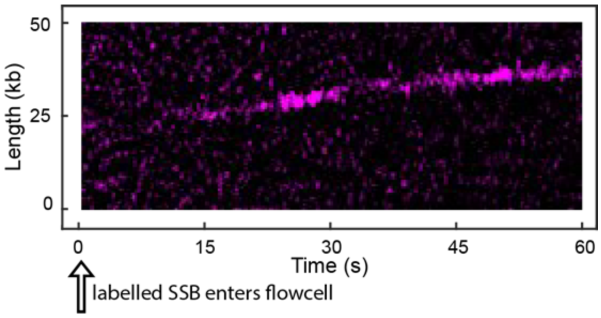


**Supplementary Figure S5.** The efficiency of internal transfer of labelled SSB and wild-type SSB are similar. The reaction was initiated and allowed to proceed for 30 s in the presence of 10 nM wild-type SSB. At *t* = 0, labelled SSB (10 nM) enters the flowcell. After ~15 s, fluorescence appears at the replication fork indicating that labelled SSB has exchanged with wild-type SSB. This exchange happens on a timescale that is similar to that shown in Figure 3E.
